# The optimal discovery procedure for significance analysis of general gene expression studies

**DOI:** 10.1101/571992

**Authors:** Andrew J. Bass, John D. Storey

## Abstract

Analysis of biological data often involves the simultaneous testing of thousands of genes. This requires two key steps: the ranking of genes and the selection of important genes based on a significance threshold. One such testing procedure, called the ‘optimal discovery procedure’ (ODP), leverages information across different tests to provide an optimal ranking of genes. This approach can lead to substantial improvements in statistical power compared to other methods. However, current applications of the ODP have only been established for simple study designs using microarray technology. Here we extend this work to the analysis of complex study designs and RNA sequencing studies. We then apply our extended framework to a static RNA sequencing study, a longitudinal and an independent sampling time-series study, and an independent sampling dose-response study. We find that our method shows improved performance compared to other testing procedures, finding more differentially expressed genes and increasing power for enrichment analysis. Thus the extended ODP enables a superior significance analysis of genomic studies. The algorithm is implemented in our freely available R package called edge.

## 1 Introduction

In genomic studies, gene expression measurements for thousands of genes are obtained simultaneously using RNA-Seq or DNA microarray technology. A primary objective in these studies is to discover biologically important genes by applying appropriate statistical tools to the data. One such approach is to apply a hypothesis testing procedure on a gene-by-gene basis to detect differentially expressed genes; for example, a *t*-test or *F*-test is commonly used to compare multiple biological groups. These test statistics are then ranked and a subset of genes with values above a specified threshold are deemed statistically significant. The significance threshold is chosen to control the false discovery rate (FDR), i.e., the proportion of false positives in the subset of selected genes. Thus there are two key steps when selecting important genes: the ranking of test statistics and the selection of tests based on a significance threshold.

One commonly used testing procedure to rank genes is the likelihood ratio test (LRT). The LRT compares the goodness-of-fit between two models, namely, the alternative and null models. The test statistic is the ratio of the likelihood under the alternative model to the likelihood under the null model, where large values indicate evidence against the null model. It is popular due to its optimality guarantees: the Neyman-Pearson (NP) lemma states that the LRT statistic is the most powerful testing procedure for a single hypothesis, i.e., no other testing procedure can achieve more power at a fixed significance threshold [1]. However, the LRT statistic may not provide the optimal ranking for multiple hypotheses [2]. This is problematic in genomics where thousands of tests are typically performed.

When there are multiple hypotheses, such as in gene expression data, many testing procedures can improve upon the statistical power by utilizing information across genes. One such method is the ‘optimal discovery procedure’ (ODP), which maximizes the number of expected true positives for a fixed number of expected false positives—a quantity related to the FDR [2]. The ODP is a generalization of the NP lemma: while the NP lemma is optimal for a *single* hypothesis test, the ODP is optimal for *italicple* hypothesis tests. The ODP achieves the optimal ranking of test statistics by leveraging information across all tests when calculating the test statistic for each gene. Intuitively, the improvement in ranking stems from functionally related genes that follow similar patterns of expression. This information is incorporated into the test statistic to strengthen or weaken the evidence for differential expression. In Storey et al. (2006), an approximation to the ODP performs favorably on DNA microarray studies compared to SAM [4], shrunken *t*-test or *F*-test [5, 6], Bayesian local FDR [7], and posterior probabilities [8].

There are two main limitations when applying the ODP to genomic studies. First, the method was primarily developed for simple static experiments (e.g., comparing two conditions) and it has not yet been extended to more complex sampling designs. Second, the underlying assumption in the ODP is that the data are generated from a Normal distribution where the per-gene observations have the same variance (i.e., homoscedasticity). This is problematic for RNA sequencing studies where the data are modeled using an over-dispersed Poisson distribution or a Normal distribution where the pergene observations have different variances (i.e., hetereoscedasticity) [9]. Due to these constraints, the applicability of the ODP has been limited to static DNA microarray studies.

In this work, we extend the ODP to both complex study designs and RNA sequencing studies. In order to incorporate dynamical responses commonly found in non-static studies, we utilize the regression spline methodology from Storey et al. (2005). A benefit of this approach is that it flexibly models gene expression responses within the ordinary least squares framework where the data are assumed to follow a Normal distribution with constant variance: this enables for a straightforward application of the ODP. For RNA sequencing studies, we implement the same strategy in ref. [9] to estimate the pergene hetereoscedasticity using the observed mean-variance relationship. We then use these estimated weights in a weighted least squares algorithm to adjust for unequal variances among observations. This transformation allows for the standard ODP framework to be utilized.

We apply our algorithm to three different experimental designs. The first is a ‘static sampling’ experiment, where samples are obtained at a fixed time point. For this example, we analyze a smoker study where smoking and non-smoking groups are compared to detect transcriptional differences in airway basal cells using RNA-Seq technology. The second is an ‘independent sampling’ experiment, where subjects are independently sampled across time or dosage-level. Here we consider two independent sampling studies, namely, a time-series and a dose-response study. The former considers the effect of age on gene expression in the cortex region of the kidney and latter explores breast cancer cell sensitivity in response to multiple 17*β*-estradiol doses. The final design is a ‘longitudinal sampling’ experiment, where subjects are sampled at multiple time points. As an example, we examine an endotoxin study which compares the leukocytes at a control group to those of an endotoxin-treated group across multiple time points.

The paper is outlined as follows. Section 2 reviews background on the ODP and regression splines. We also review a computationally efficient implementation of the ODP called the ‘modular optimal discovery procedure’ (mODP). Section 3 introduces our proposed algorithm and Section 4 illustrates the results from our method on the four studies. We validate these results through comprehensive simulations.

## 2 Background

### 2.1 The optimal discovery procedure

The optimal (or ‘most powerful’) hypothesis testing algorithm for a *single* test is provided by the Neyman-Pearson (NP) lemma [1]. Given some observed data **y** = (*y*_1_, *y*_2_, *…, y*_*n*_), the NP lemma states that the ratio of the alternative likelihood *g*(**y**) over the null likelihood *f* (**y**)—known as the likelihood ratio— has the largest power for each false positive rate compared to any other decision rule. Intuitively, this optimality arises because the data generating process under each model is assumed to be known. For multiple hypotheses, the likelihood ratio is applied on a case-by-case basis. However, potentially useful information across different hypotheses are ignored. As a consequence, the likelihood ratio may no longer be an optimal decision rule [2].

The ‘optimal discovery procedure’ (ODP) is a generalization of the NP lemma for multiple hypotheses. More specifically, consider gene expression measurements (y_1_, y_2_, …, y_*m*_) where there are *m* genes and *n* samples. Further assume that the first *m*_0_ and the last *m*_1_ = *m − m*_0_ hypotheses are from the alternative and null models, respectively. The ODP test statistic for gene *i* is

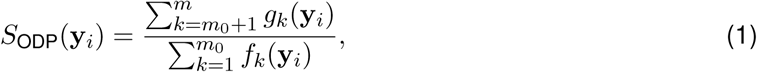

where *g*_*k*_(**y**_*i*_) is alternative likelihood and *f*_*k*_(**y**_*i*_) is the null likelihood under gene *k*. The numerator and denominator can be viewed as the cumulative power under the alternative and null models, respectively, across all hypotheses: only tests related to gene *i* contribute to the above statistic. Storey (2007) shows that this test statistic maximizes the number of expected true positives (ETP) for a fixed number of expected false positives (EFP)—a quantity closely related to the false discovery rate, i.e., EFP+ETP 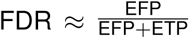 Therefore, by leveraging information across different hypotheses, the ODP achieves the optimal *ordering* (or ranking) of test statistics. Moreover, the improvements in statistical power are generally substantial compared to the likelihood ratio test (see [2]).

Evaluating Equation (1) requires making assumptions on the data generating process and the hypothesis status for each test. In this work, the alternative and null densities follow a Normal distribution with some finite mean and variance. However, it is not known *a priori* which tests are from the alternative and null models. Instead an approximation to the true ODP statistic is estimated [3], i.e. 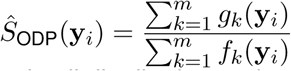. Another complication is that the ODP test statistic does not have a theoretical null distribution and so *p*-values cannot be analytically calculated. Therefore, a bootstrap procedure must be implemented to generate an empirical null distribution of the test statistics. This estimated ODP has been shown to provide similar power to the true ODP [3]. However, calculating the test statistics involves 2*m*^2^ computations and so it is computationally demanding for genomic datasets where *m* can range anywhere from 10^3^ to 10^5^.

#### Algorithm 1 KL clustering algorithm for the modular optimal discovery procedure (mODP)

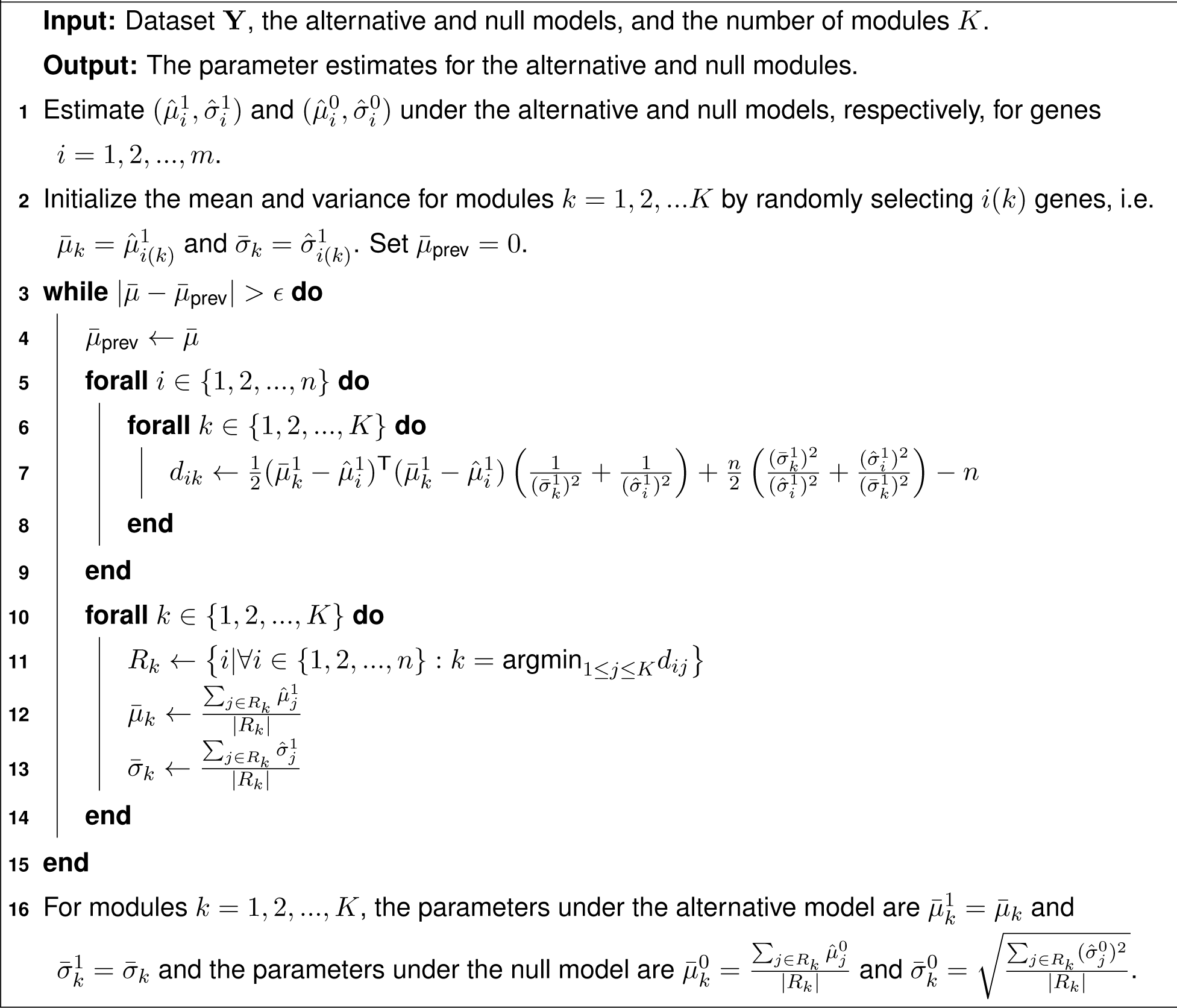

Woo et al. (2010) proposed a computationally efficient approximation to the ODP called the ‘modular optimal discovery procedure’ (mODP). Similar to the estimated ODP, the mODP assumes that the data are generated from a Normal density with parameters 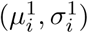 and 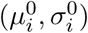 for genes *i* = 1, 2, *…, n* under the alternative and null models, respectively. These parameters are estimated from the data using an ordinary least squares algorithm. A clustering algorithm then assigns genes to *k* = 1, 2, *…, K* modules based on the Kullback-Leibler distance *d*_*ik*_ (only the alternative model is used to determine gene-module assignments). Using the module assignments, the parameters are updated and genes are reassigned to new modules: the above steps continue until a convergence criteria is met. The final module parameters are denoted by 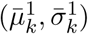 and 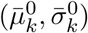 under the alternative and null models, respectively (see Algorithm 1).

Given the module parameters, the mODP test statistic can be expressed as

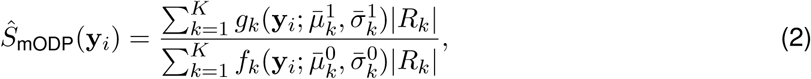

where 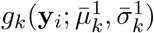 is the alternative likelihood and 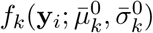 is the null likelihood under module *k*, and *|R*_*k*_*|* is the number of genes belonging to module *k*. A bootstrap algorithm is implemented to generate the empirical null distribution of the test statistics (described in Appendix 7.2). The mODP reduces the number of calculations from 2*m*^2^ to 2*Km* where *K* ≪ *m*. Thus the time complexity of the mODP is linear with the number of genes. In Woo et al. (2010), the authors demonstrate that the mODP has similar power to the estimated ODP and is robust to the number of modules when *K* ≥ 50.

### 2.2 Regression splines

The general framework for modeling non-linear responses in complex study designs follows from Storey (2005): Consider an experiment with gene expression measurements *y*_*ij*_ and explanatory variable *x*_*j*_ for *i* = 1, 2, *…, M* genes and *j* = 1, 2, *…, N* samples. In non-static studies, there can be multiple measurements of *x*_*j*_ for sample *j*, i.e., *x*_*jk*_ where *k* = 1, 2, *…, T*_*j*_; for example, *x*_*jk*_ can be multiple time points or different dosage levels for a particular sample. The expression for gene *i* is modeled as

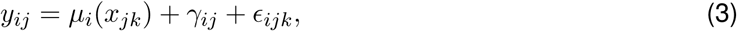

where *µ*_*i*_(*x*) is the population average curve, γ_*ij*_ is the individual-specific random deviation from the population average curve, and ε_*ijk*_ is a random error that follows a Normal distribution with mean 0 and variance 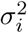. Here we assume that the individual-specific random effects follow a Normal distribution with mean 0 and constant variance 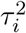.

The population average curve can be flexibly modeled using a regression spline. A regression spline is a piecewise polynomial function continuous at *d* specified points (or ‘knots’). We only consider natural cubic splines, which are third order polynomial functions that are linear beyond the boundary knots. In this case, the population average curve can be parameterized by a *d*-dimensional basis, i.e. 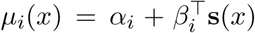 where s(*x*) = (*s*_1_(*x*), *s*_2_(*x*), *…, s*_*d*_(*x*)) is a prespecified *d*-dimensional natural cubic spline basis and the parameters *α*_*i*_ and *β*_*i*_ = (*β*_*i*1_, *β*_*i*2_, *…, β*_*id*_) are estimated by ordinary least squares. The parametric model for *µ*_*i*_(*x*) enables testing of parameters *α*_*i*_ and *β*_*i*_, which do not depend on specific *x*: this simplification allows for inferences of general sampling designs [10]. We apply this framework to a static sampling study, two independent sampling studies, and a longitudinal sampling study.

In a static sampling study, subjects are independently sampled across one or more biological groups at a fixed time point. As an example, in the smoker study *x*_*j*_ is a categorical variable indicating the smoking status of individual *j* and *y*_*ij*_ is the RNA-Seq count for gene *i* (Figure 1a). Equation (3) can be simplified by modeling the population average curve as *µ*_*i*_(*x*_*j*_) = *α*_*i*_ + *β*_*i*_*x*_*j*_. In this study, we are interested in determining whether gene expression is differentially expressed between groups (the alternative hypothesis) or remains unchanged (the null hypothesis). Therefore, the null hypothesis model (dashed line) is fit under the constraint of *β*_*i*_ = 0 and the alternative hypothesis model (solid line) allows this parameter to be unconstrained.

**Figure 1:**
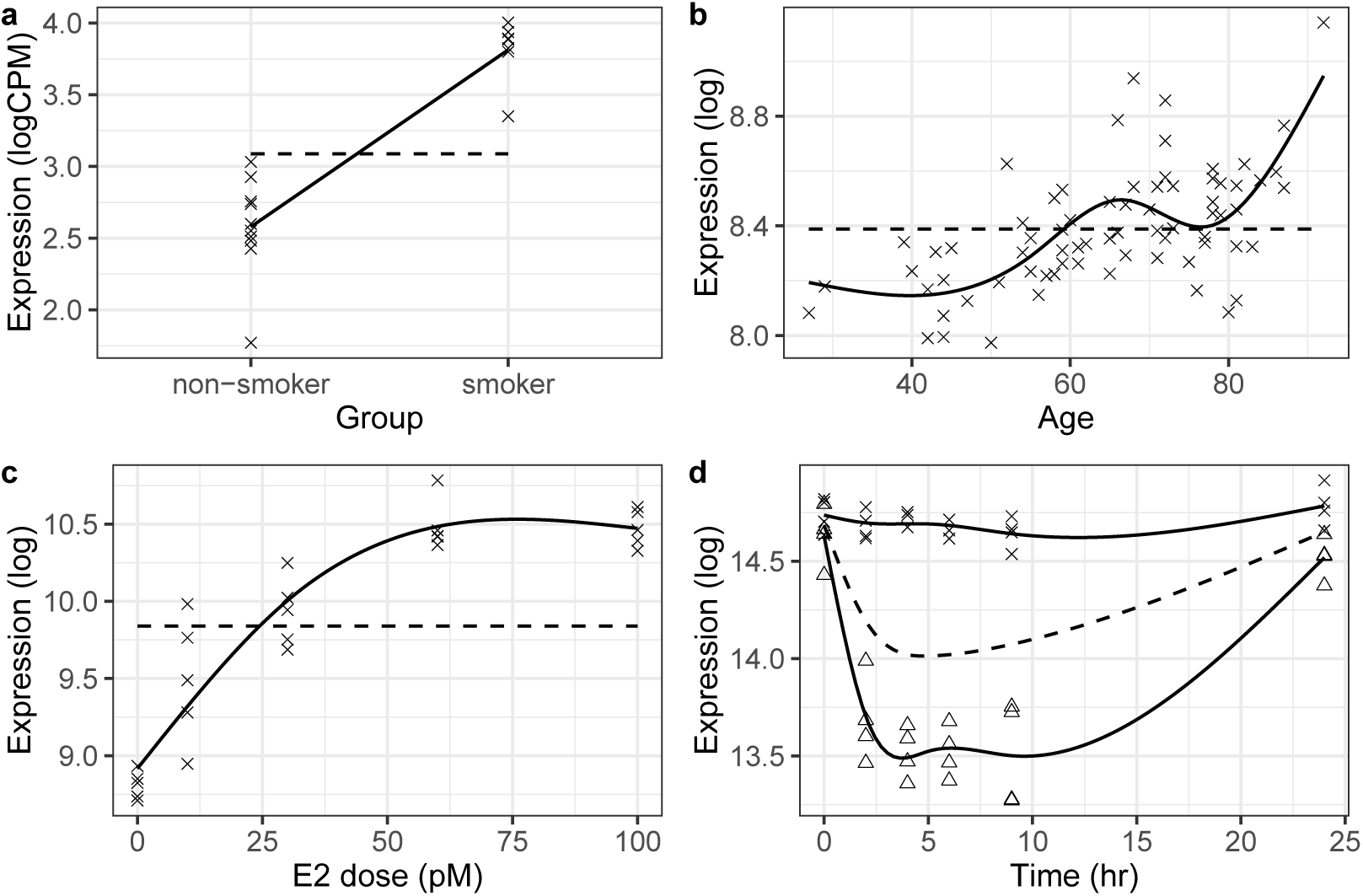
Fitting regression splines to general study designs: (a) static, (b-c) independent sampling, and (d) longitudinal studies. The null (dashed) and alternative (solid) models are shown for a significant gene. In (d), the endotoxin-treated and control groups are denoted by a triangle and cross, respectively.

In an independent sampling study, subjects are independently sampled across a continuous variable (similar to cross-sectional sampling). The population average curve is modeled using a *d*-dimensional natural cubic spline basis. There are two studies analyzed with independent sampling designs. The first is a kidney aging study where human subjects are independently sampled at various ages (Figure 1b). The second is a dose-response study where 17*β*-estradiol is introduced to breast cancer cells at various dosage levels (Figure 1c). In both of these studies, the objective is to determine whether gene expression is differentially expressed across time or dosage level (the alternative hypothesis) or remains unchanged (the null hypothesis). Therefore, the null hypothesis model (dashed line) is fit under the constraint of *β*_*il*_ = 0 for *l* = 1, 2, *…, d* and the alternative hypothesis model (solid line) allows these parameters to be unconstrained.

#### Algorithm 2 Algorithm for modeling non-linear responses

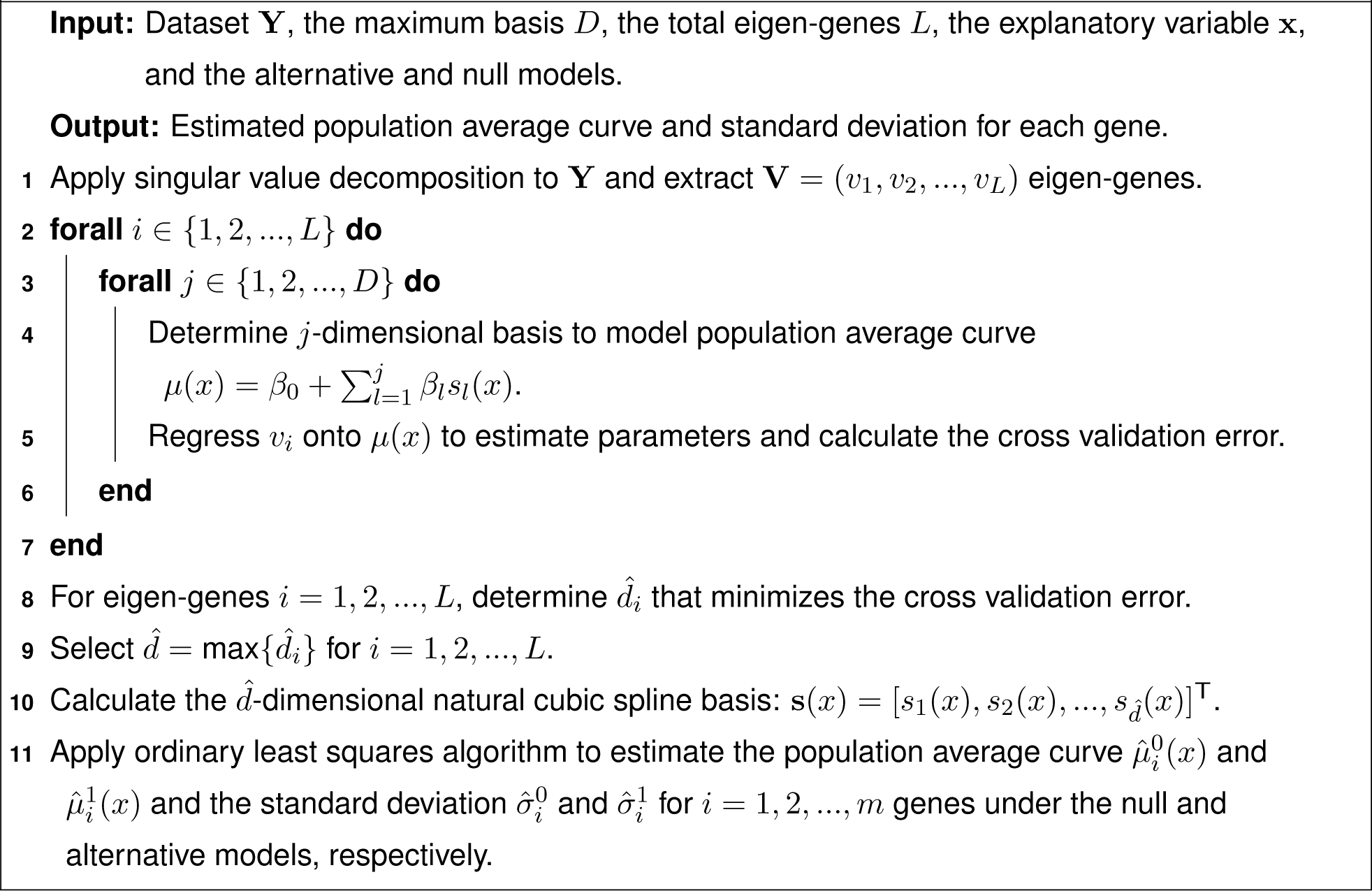

In a longitudinal sampling study, subjects are sampled multiple times across a continuous variable. The response variable can be modeled using Equation (3) where the population average curve is a *d*-dimensional natural cubic spline basis. As an example, the endotoxin study compares two different classes across time, namely, endotoxin-treated versus control-treated. For this case, *y*_*ij*_ is the gene expression measurement for gene *i* in individual *j* and *x*_*jk*_ indicates the time point individuals were sampled (Figure 1d). The alternative hypothesis is that there is differential expression between classes while the null hypothesis is that there is no difference in gene expression. Thus the null hypothesis model fits one curve to both classes combined (dashed line) and the alternative hypothesis model fits two separate curves to each class (solid line).

Algorithm 2 summarizes the model fitting procedure to estimate the population average curve in a study. While natural cubic splines provide flexible parametric models, they require the placement of *d* knots. We utilize the cross validation algorithm in ref. [10] to automatically choose the optimal 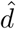. First, we apply a singular value decomposition to extract the top *i* = 1, 2, *…, L* eigen-genes. We then regress these eigen-genes onto s(*x*) = (*s*_1_(*x*), *s*_2_(*x*), *…, s*_*j*_(*x*)), where we use a *j* = 1, 2, *…, D* dimensional natural cubic spline basis: the knots are placed at evenly spaced quantiles, i.e. the 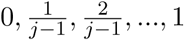 quantiles. Finally, the basis dimension used for model fitting is chosen by applying a cross validation procedure to select the optimal 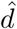 across all eigen-genes. Using the selected 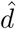, an ordinary least squares algorithm is applied to estimate the population average curve and variance for all genes under the alternative and null models.

## 3 Methods

Our algorithm introduces regression splines into the ODP framework to extend it to complex study designs. To do so, the ODP test statistic must be extended to incorporate non-linear responses. Suppose the data are **y**_*i*_ and the explanatory variable is *x*_*j*_ where there are *i* = 1, 2, *…, m* genes and *j* = 1, 2, *…, n* samples. In either case, there are two different models, namely, the null model with parameters 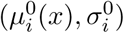 and the alternative model with parameters 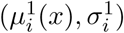. The objective is to test the null hypothesis 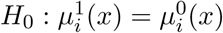 versus the alternative hypothesis 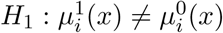. In this work, the population average curves are flexibly modeled using a *d*-dimensional natural cubic spline basis. The parameters under both models can then be estimated by ordinary least squares, i.e., 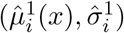 and 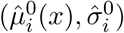

For non-linear responses, the estimated optimal discovery procedure is

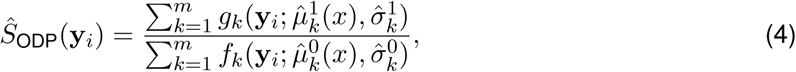

where the likelihoods are assumed to follow a Normal distribution. It is evident that 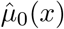 is not of interest in the testing procedure (so-called ‘nuisance parameter’): this ancillary information can be removed by transforming the data to 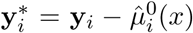. Under this transformation, the hypotheses are 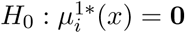 versus 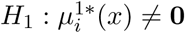. This modified version of ODP is

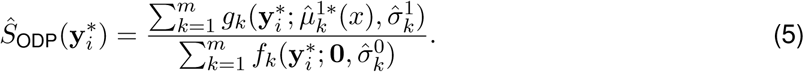

Similar to the original implementation of the ODP, the above test statistic requires 2*m*^2^ calculations which makes it computationally slow for genomic data sets. Instead, we can incorporate the population average curve into the mODP as

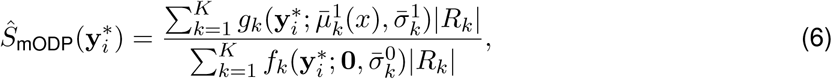

where there are *k* = 1, 2, *…, K* modules, the membership size of module *k* is *|R*_*k*_*|*, and the module parameters are estimated by applying the mODP clustering algorithm (described in Algorithm 1).

### Algorithm 3 Algorithm for analyzing complex study designs

**Input:** Dataset **Y**, the alternative and null model, the number of bootstrap iterations *b*, and the number of modules *K*.

**Output:** Empirical *p*-values for each gene.

**1** Apply Algorithm 2 to estimate the population average curves and standard deviations.

**2** Subtract the null model from the observed data, 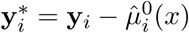, and estimate 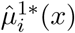 and 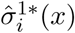 for *i* = 1, 2, *…, m* genes

**3** Estimate the module parameters under the null and alternative models using Algorithm 1 and calculate 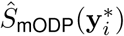 for *i* = 1, 2, *…, m* genes.

**4** Generate *k* = 1, 2, *…, b* datasets from the null model by resampling the scaled residuals 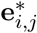 from the alternative model fit and add it to 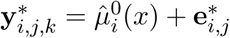 (described in Appendix 7.2).

**5** Apply (2-3) to generate the null test statistics 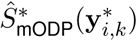 for *k* = 1, 2, *…b* bootstrap iterations and *i* = 1, 2, *…, m* genes.

**6** Using the observed and null test statistics, calculate empirical *p*-values.

Our proposed method is summarized in Algorithm 3: The inputs are the observed dataset **Y**, the alternative and null models, the number of modules *K* and the number of bootstrap iterations *b*. First, we apply Algorithm 2 to explanatory variable **x** to model the non-linear gene expression responses. The data are then transformed by subtracting the null model fit from the observed dataset. This adjusted dataset 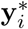 is regressed onto the alternative model to get updated parameter estimates, 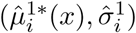. Finally, we apply the mODP clustering algorithm to determine the parameters for the *k* = 1, 2, *…, K* modules under the alternative and null models, 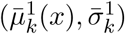 and, 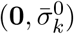 respectively (see Algorithm 1). Using the parameter estimates from the clustering algorithm, the mODP statistic is calculated for all genes. A bootstrap algorithm is implemented to calculate the empirical null distribution of the test statistics (described in Appendix 7.2). For the datasets in this paper, there are *b* = 500 bootstrap iterations (*b* = 5000 for the smoker study) and *K* = 800 modules.

Prior applications of ODP are focused on microarray studies where it is common to assume that the data generating process is approximately Normal and homoscedastic. However in sequencing studies, this assumption is no longer valid because the observations are heteroscedastic. In order to apply the ODP to sequencing studies, we implemented a similar strategy in ref. [9] where RNA-Seq data are log-transformed to model the observed mean-variance relationship. Using this model, weights capturing the heteroscedasticity across observations are estimated. These weights are then incorporated in a weighted least squares regression and are easily integrated into the mODP framework: Given a set of variance stabilizing weights *w*_*ij*_ and a *d*-dimensional natural cubic spline basis **s**(*x*_*ij*_) for *j* = 1, 2, *…, n* samples and *i* = 1, 2, 3*…, m* genes, the data are transformed as s^*\**^(*x*_*ij*_) = *w*_*ij*_s(*x*_*ij*_) and 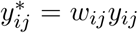 An ordinary least squares algorithm can then be applied to this transformed data. Thus Algorithm 3 can be appropriately adjusted to accommodate these weights.

## 4 Results

The modular optimal discovery procedure (mODP) is applied to four different genomic experiments. The performance of mODP is compared to an *F*-test and a moderated *F*-test [6] using the number of discoveries and enriched gene sets. Finally, we validate our findings through comprehensive simulations.

### 4.1 Datasets

#### Kidney study

To elucidate the transcriptional response from aging in the kidney, the kidney study collected cortex samples from 72 patients with ages ranging from 27 to 92 years [11]. The samples were hybridized onto U133a and U133b GeneChips with 44,928 probes. Following similar filtering steps in Storey (2005) to control for potential confounding, only 38,833 probes were used for analysis and the expression values were log-transformed for variance stabilization.

#### Endotoxin study

The endotoxin study analyzed transcriptional regulation in human blood leukocytes from two experimental groups: a treatment group receiving a bacterial endotoxin (an inflammatory stimulus) and a control group [12]. There were four samples in each biological group and blood samples were collected at 2, 4, 6, 9, and 24 h intervals. One control sample had missing information at the 4 and 6 hr time points. The samples were hybridized onto U133 GeneChips with 44,924 probes. The expression values were log-normalized for variance stabilization.

#### Smoker study

The smoker study is a two group comparison between smoking and non-smoking humans [13]. There are a total of 17 samples (10 non-smokers and 7 smokers) from human airway basal cells in the epithelium: there is one female smoker and the rest of the samples are males. The samples are sequenced (paired-end) using Illumina HiSeq 2000 and the reads are assembled using Bowtie: there are total of 65,217 genes with mapped reads. After filtering genes with fewer than 10 reads across all samples, only 26,268 genes remained for analysis. The R package limma is used to estimate the inverse-variance weights for the weighted least squares implementation. The expression values were transformed to log2-counts per million (logCPM).

#### Dose study

The dose study is a dose-response experiment where sensitivity to 17*β*-estradiol (E2) in breast cancer cells (BUS cells) was examined [14]. There are five biological replicates for each E2 concentration, where the E2 concentrations considered were 0, 10, 30, 60, and 100 pM (25 total samples). After 48 hours exposed to E2, RNA samples were hybridized onto U133a GeneChips with 22,283 probes. The expression values were log-normalized to stabilize the variance.

### 4.2 Determining the degrees of freedom

We implemented the cross validation procedure detailed in Algorithm 2 to determine the appropriate degrees of freedom *d* for the natural cubic spline basis. (The smoker study is a two group comparison and so regression splines are not necessary.) For each study, the first four eigen-genes were determined by applying a singular value decomposition to the dataset. In the endotoxin study, the control-treated and endotoxin-treated groups were separated into two distinct datasets. Multiple regressions were fit to the eigen-genes using *d* = 1, 2, 3, 4 for the endotoxin and dose studies and *d* = 1, 2, …, 10 for the kidney study (an intercept term was included in the model). For each eigen-gene, the *d* that minimized the leave-one-out cross validation error was selected. Finally, the maximum *d* across all eigen-genes was chosen as the estimated degrees of freedom. Applying the above procedure, we find 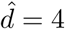 for the endotoxin and kidney studies and 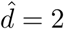 for the dose study (Figure 2).

**Figure 2:**
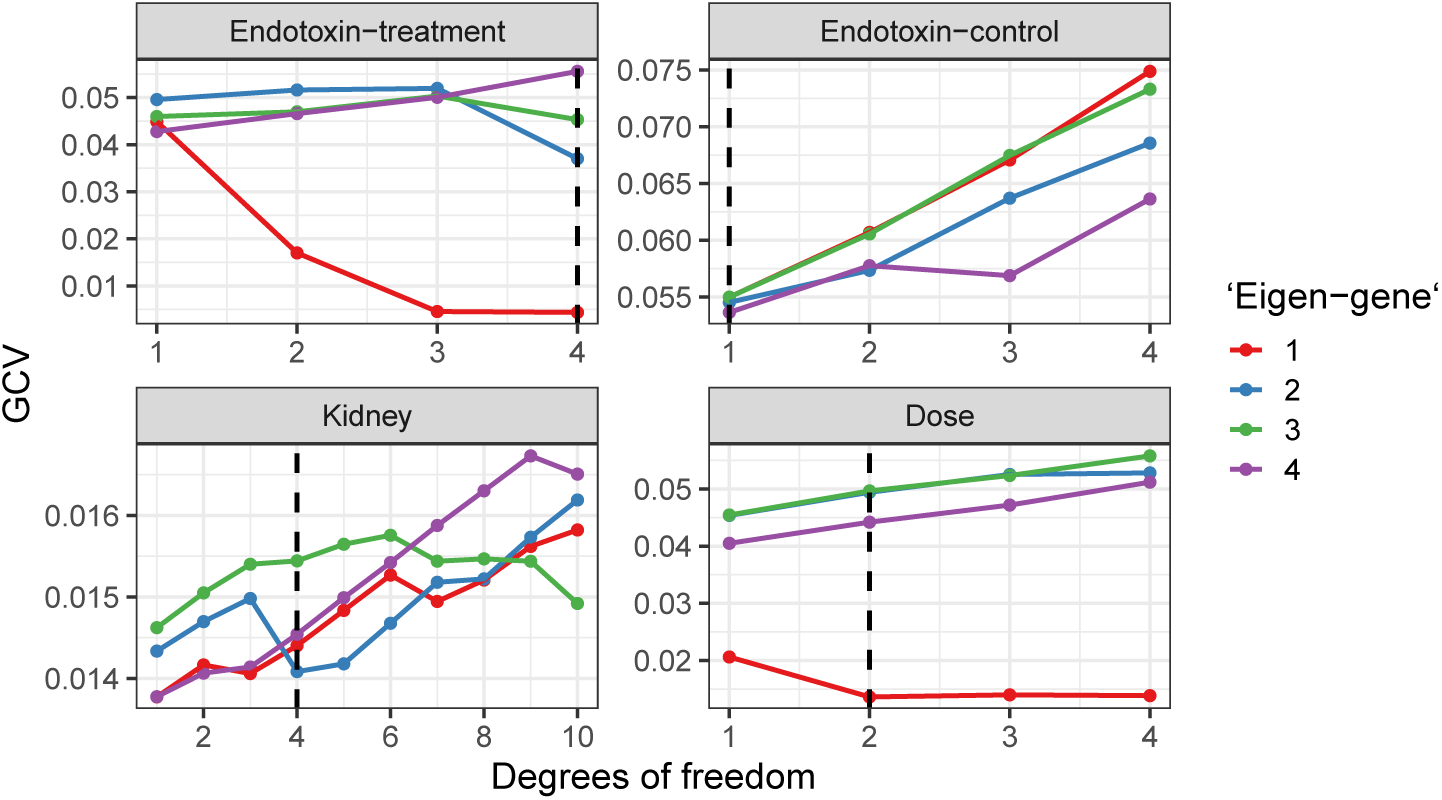
Cross validation error versus degrees of freedom for the first four eigen-genes. The dotted line indicates the chosen degrees of the freedom in the study.

### 4.3 Method comparisons

We compared the mODP to two other popular test statistics, namely, the *F*-statistic and the moderated *F*-statistic (described in Appendix 7.1). Compared to the *F*-statistic, the moderated *F*-statistic shrinks the sample variance towards a pooled variance. This shrinkage allows for more stable inferences with low sample size studies [6]. Unlike the mODP which requires an empirical null distribution, the *F*-statistic and moderated *F*-statistic have theoretical null distributions. Therefore, we also estimate an empirical null distribution for the *F*-test and moderated *F*-test using a bootstrap algorithm (described in Appendix 7.2). In summary, the mODP is compared to an *F*-test, a moderated *F*-test, a bootstrap *F*-test and a bootstrap moderated *F*-test.

We applied the above testing procedures to our four chosen studies and calculated the number of differentially expressed genes at various false discovery rates (Figure 3). At each false discovery rate threshold, the mODP finds substantially more differentially expressed genes compared to the other methods. For example, when applying a false discovery rate of 0.1, mODP detects 1481, 297, 6637, and 887 more differentially expressed genes in the kidney, smoker, endotoxin and dose studies, respectively. Additionally, the mODP finds nearly all of the differentially expressed genes detected by the other methods (Figure 4). Finally, we find that the mODP has the lowest estimated proportion of true nulls across all studies (Table 1). This suggests that the mODP estimates a higher expected number of alternative genes.

**Figure 3:**
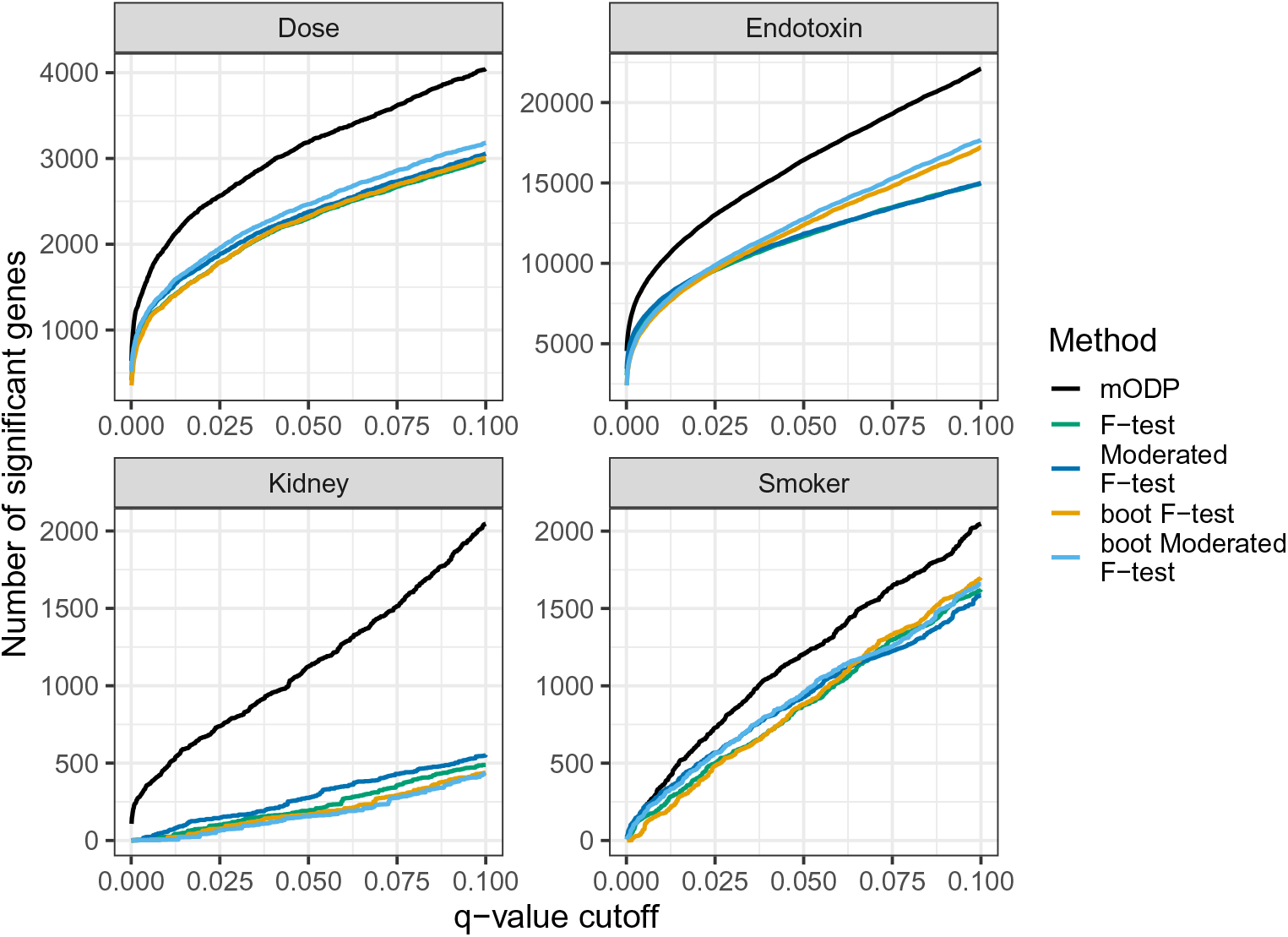
Observed number of discoveries at various *q*-value cutoffs from the modular optimal discovery procedure (black), F-test (green), bootstrap F-test (orange), moderated F-test (blue) and bootstrap moderated F-test (light blue). These methods were applied to four different studies: endotoxin (longitudinal sampling), kidney (independent sampling), dose (independent sampling), and smoker (static sampling).

**Figure 4:**
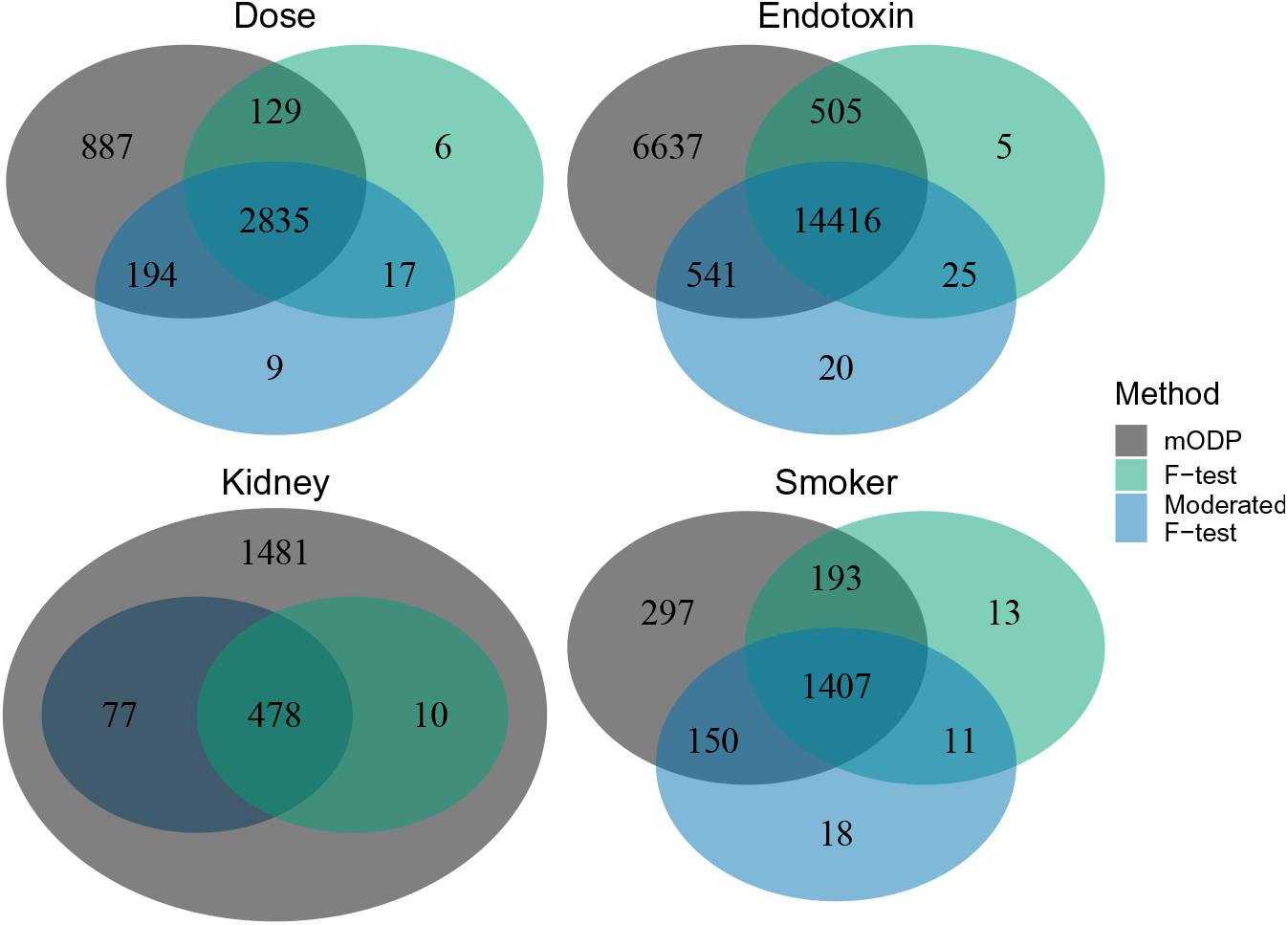
Venn diagram of the total discoveries at a false discovery rate of 0.1 for the studies in Figure 3. Only the modular optimal discovery procedure (black), F-test (green) and moderated F-test (blue) are shown.

**Table 1:** Estimated proportion of true nulls.

| | ODP | $F$-test | Mod. $F$-test | boot. $F$-test | boot. mod. $F$-test |
| --- | --- | --- | --- | --- | --- |
| Dose | 0.672 | 0.803 | 0.813 | 0.804 | 0.793 |
| Endotoxin | 0.349 | 0.605 | 0.615 | 0.388 | 0.397 |
| Kidney | 0.585 | 0.687 | 0.685 | 0.684 | 0.676 |
| Smoker | 0.597 | 0.681 | 0.681 | 0.680 | 0.675 |

To compare the testing procedures, we also performed an enrichment analysis using the hallmark gene sets from the MSigDB database. These gene sets contain highly curated genes with clear expression for well-defined biological states or processes [15, 16]. We developed a simple procedure to detect important gene sets by assigning the proportion of true positives to each. These values range from 0 to 1, with important gene sets having largest values (see Appendix 7.4 for additional details). We find that the mODP has the largest proportion of true positives across all gene sets compared to other methods (Figure 5). Thus the mODP has more power to detect gene sets with enriched true positives.

**Figure 5:**
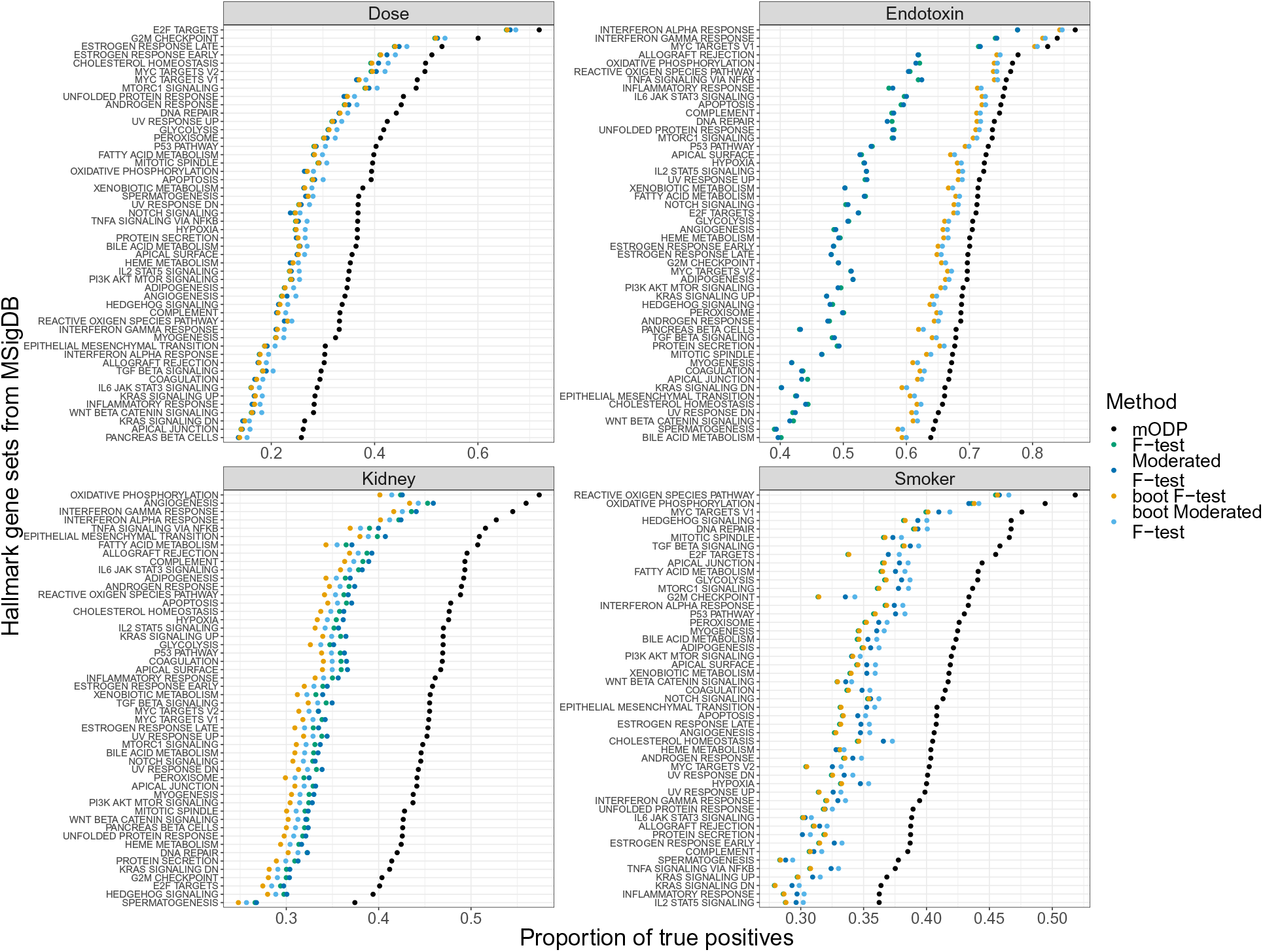
Proportion of true positives for the Hallmark gene sets from MSigDB. Enrichment results for each method are shown for the dose, endotoxin, kidney, and smoker studies.

### 4.4 Simulations

Comprehensive simulations were performed to verify the observed differences between mODP and other methods. We generated 500 representative datasets of the observed studies as follows. For each study, the *F*-testing procedure was used to separate genes into two distinct classes (i.e., alternative and null) based on a false discovery rate threshold of 0.1. We then sampled from the population of alternative genes to get unique gene expression profiles. In total, we considered 5, 10, 50, 100, and 200 unique gene expression curves in our simulation studies. These curves defined the signal for the alternative genes. (Note the smoker study is a static experiment and so ‘unique gene expression profile’ refers to the mean differences between the two conditions.) The signals for the null genes were sampled from the null population. Random noise was added to maintain the signal-to-noise ratio and to match the observed power. Finally, the total number of alternative and null genes were chosen to keep the observed proportion of true nulls fixed. For more details see Appendix 7.3.

To compare the testing procedures in the simulated datasets, we calculated the estimated false discovery rate (FDR) and the total number of discoveries. We find that the mODP controls the FDR in all simulated studies (Figure 6b). Furthermore, substantially more differentially expressed genes were detected relative to the other testing procedures (Figure 6a). We also find that the moderated *F*-test identifies a similar number of differentially expressed genes compared to the *F*-test. This is unsurprising as the moderated *F*-test only outperforms the *F*-test when there are very small sample sizes. When the number of unique gene expression patterns is increased, the power of the mODP decreases. This is because the number of unique gene expression curves does not change the power of the *F*-test and moderated *F*-test.

**Figure 6:**
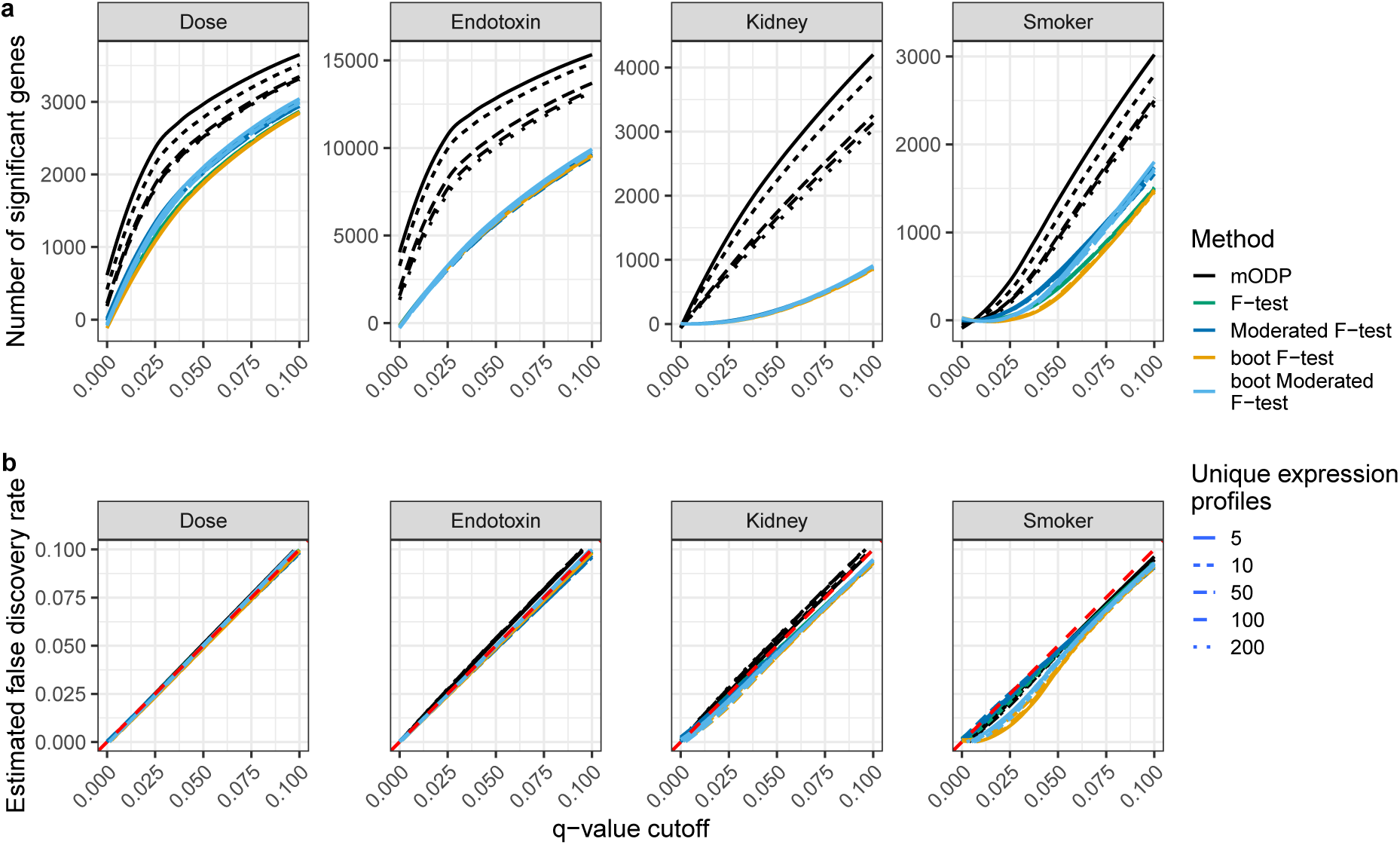
Simulation results for the dose, endotoxin, kidney, and smoker studies using 5, 10, 50, 100, and 200 unique gene expression profiles (linetype). The modular optimal discovery procedure (black), F-test (green), moderated F-test (blue), bootstrap F-test (orange) and bootstrap moderated F-test (light blue) are applied to the simulated studies. **(a)** Estimated power at multiple *q*-value cutoffs between 0.0001 and 0.1. **(b)** Estimated false discovery rate. Curves represent the average value from 500 replications.

## 5 Discussion

The ODP is a test statistic that provides substantial improvements in statistical power compared to other testing procedures. While previous work on the ODP is limited to static microarray studies [3, 2, 17], here we extend its application to complex experimental designs and sequencing studies. Our proposed algorithm is applied to two time series studies, a dose-response study and an RNA-Seq study. For each study, our method detects more differentially expressed genes and improves the statistical power for gene set enrichment analysis. These improvements in power are validated through comprehensive simulations, where data are simulated to closely resemble the observed datasets. Therefore, the ODP allows for a more thorough investigation of underlying biological mechanisms in downstream analysis.

The gained improvements in power from the ODP have important biological implications. In a genomic study, genes share similar patterns of gene expression based on their functional roles. The ODP leverages this information across genes to strengthen the evidence for or against differential expression. To explore how the ODP depends on the ‘degree’ of functionally related genes, we varied the number of unique gene expression profiles with simulated data. We find that the ODP loses power as the number of unique gene expression profiles increases: there are fewer functionally related genes and so there is less information that can be leveraged in the test statistic. As an extreme case, suppose all genes in a study are functionally unrelated. In this scenario, the ODP has been shown to perform similar to the *F*-test (see [3]). While this example is unlikely in biological studies, it provides intuition for the observed power improvements compared to other testing procedures.

There are a few considerations to note when applying our extended framework to genomic datasets. First, the computationally efficient implementation of the ODP, called the modular ODP (mODP), involves specifying the number of modules. While previous work has recommended 50 modules for static microarray studies, we found choosing at least 200 modules to capture the complex functional relationships among genes provides the best results. Second, there needs to be an adequate number of samples in the study. This is due to the constraints of the mODP: it requires accurate estimates of the mean and variance. Furthermore, the bootstrap algorithm implemented in the procedure requires a minimum number of samples per biological condition to generate a valid empirical null. In the studies considered here, there are at least four biological replicates per condition. Lastly, the appropriate degrees of freedom needs to be carefully chosen to avoid overfitting the spline. To this end, we implemented a procedure from ref. [10] that chooses the degrees of freedom based on the leave-one-out cross validation algorithm.

An interesting aspect of the mODP implementation is that a clustering algorithm assigns genes to modules, where the modules are representative of shared functional gene expression patterns. These modules provide valuable information that can be utilized in an exploratory data analysis. For example, we can calculate the proportion of true positives for each module and rank modules based on true positive enrichment. Modules enriched with true positives can then be further analyzed to understand functional relationships among genes. Thus the clustering algorithm provides information of potential use in other biological analyses.

There are a number of ways the ODP can be further extended for genomic studies. One enhancement is incorporating prior weights on each hypothesis test. For example in sequencing data, higher per-gene read counts are more reliable than lower per-gene read counts. This information can be included into a weighted ODP, where weights are generated by estimating the functional proportion of true nulls based on some informative variable [18]. Another improvement is to extend the ODP to generalized linear models where the response variable follows an exponential family distribution. This would enable its extension to genome-wide association studies where complex traits are commonly non-Normal. These avenues will be explored in future work.

As the cost of generating biological samples decreases, the prevalence of complex study designs will increase. The key motivation behind these studies is to capture inherently non-linear transcriptional responses. Therefore, there is demand for statistically rigorous methodologies that can be applied to such settings. In this work, we develop a framework to model non-linear gene expression responses while optimally utilizing biological correlations among genes to improve statistical power. Our method can thus uncover important biological insights across a wide range of applications in functional, translational, and clinical genomics.

## 6 Software and Data

An implementation of the algorithm described in this paper is available as an R package called edge. The package can be downloaded at https://github.com/StoreyLab/edge (most recent version) or https://bioconductor.org/packages/release/bioc/html/edge.html (release). The data and code used to produce the figures in this manuscript can be found at https://github.com/StoreyLab/odp_general_studies.

## 7 Appendix

### 7.1 The *F*-statistic and moderated *F*-statistic

Suppose gene expression data *y*_*ij*_ and explanatory variable *x*_*lj*_ are observed for *i* = 1, 2, *…, M* genes, *j* = 1, 2, *…, n* samples, and *l* = 1, 2, *…, d* explanatory variables; the design matrix **X** = (**x**_1_, **x**_2_, …, **x**_*d*_) is assumed to be full rank. In this case, the null model 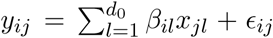 is tested versus the alternative model 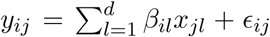 where 1 *≤ d*_0_ *< d* and ϵ_*ij*_ are uncorrelated random errors that follow a Normal distribution with mean zero and variance 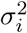. We are interested in comparing both models to infer whether *β*_*il*_ *≠* 0 for at least one of the *l* = *d*_0_ + 1, *…, d* explanatory variables.

The *F*-test is a classical testing procedure that can be used to compare nested regression models.

The general procedure works as follows. The alternative model is fit using ordinary least squares to the observed data to estimate the parameters 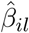 and the residual vector 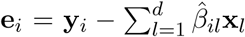. Similarly, the null model is fit to estimate the parameters 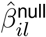 and the residual vector 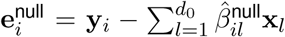 The test statistic is defined as

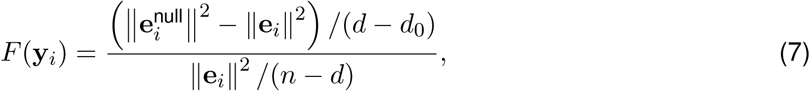

where the theoretical distribution under the null hypothesis follows Fisher’s *F*-distribution with *d - d*_0_ degrees of freedom in the numerator and *n - d* degrees of freedom denominator (denoted 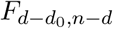). Intuitively, if there is no difference between both models then *F* (**y**_*i*_) should be concentrated around 1. Otherwise, large deviations from 1 provide evidence against the null model. The assumption that the F-statistic follows an 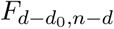-distribution under the null hypothesis is only true asymptotically: in practice, there needs to be a large number of samples for reliable inferences.

For small sample sizes, the moderated *F*-statistic can be used to compare two models. The main issue with small sample sizes is that the sample variance can often be inflated and unreliable to use in the traditional *F*-test. The moderated *F*-test is an empirical Bayes procedure that borrows information across genes to provide stable estimates of the sample variance [6]. A rough outline of the hierarchical model is as follows. The inverse variance across genes are assumed to vary as a scaled chi-squared distribution, i.e.,

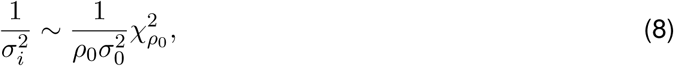

where *ρ*_0_ is the degrees of freedom and 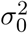 is a scaling factor. Furthermore, the non-zero effect sizes are assumed to follow a Normal distribution with mean 0 and variance proportional to 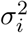. The posterior mean of 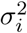 given the sample variance can be determined from the above hierarchical model, see [6] for more details. This mean value is used as an improved estimate of the sample variances, where the sample variances are shrunken towards the prior estimator 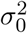 for more stable estimates. More specifically, the moderated *F*-statistic is defined as

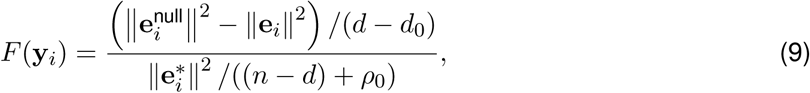

where 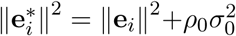 and the parameters 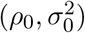 are estimated from the data [6]. The theoretical distribution under the null hypothesis for the moderated *F*-statistic follows an *F*-distribution with *d* − *d*_0_ degrees of freedom in the numerator and (*n* − *d*) + *ρ*_0_ degrees of freedom in the denominator, 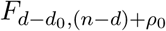. When there are a sufficient number of samples, 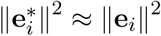 and (*n-d*)+*ρ*_0_ *≈ n-d*. Thus the statistical power of the moderated *F*-test and *F*-test will be similar for large sample sizes.

### 7.2 Generating an empirical null

A standard bootstrap procedure was implemented to generate an empirical null distribution for the testing procedures. For *i* = 1, 2, *…, M* genes,

1. Assume the model fit for the null model is **y**_*i*_ = *f*_0_(**x**_*i*_) + ϵ_*i*_ and for the alternative model is **y**_*i*_ = *f*_1_(**x**_*i*_) + ϵ_*i*_ where ϵ_*i*_ are the random errors.
2. Fit both models to the observed data using ordinary least squares: Estimate 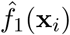 and **e**_*i*_ for the alternative model and 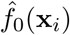 and 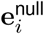 for the null model. Calculate the test statistic of interest (i.e., the ODP statistic, *F*-statistic, or moderated *F*-statistic), denoted by 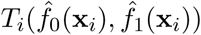
3. Adjust the residuals by calculating studentized residuals: 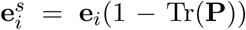, where **P** = **X**(**X**^*T*^ **X**)^*-*1^**X**^*T*^ is the projection matrix under the alternative model.
4. For *b* = 1, 2, *…, B* bootstrap samples, sample *n* observations from the studentized residuals (with replacement) to obtain 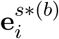. Add these residuals to the null model fit 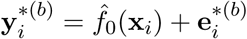.
5. Fit both models to 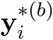 and obtain 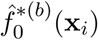 and 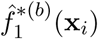 estimates under the null and alternative models, respectively. Calculate 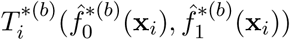 for *b* = 1, 2, *…, B* bootstrap samples. Note that the hyperparameters of the moderated *F*-statistic (*d*_0_, *σ*_0_) are fixed and so 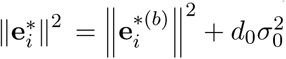
6. Calculate the empirical *p*-values as

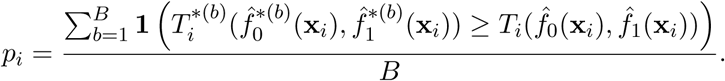

For the ODP, the studentized residuals need to be rescaled by the observed sample variance. This enforces that the sample variance remains the same for all bootstrap iterations. Thus step (4) is 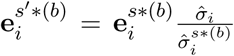 where 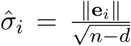 is the sample standard deviation of the residuals from the original alternative model (*d* explanatory variables) and 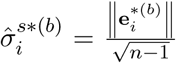 is the standard deviation from the resampled residuals.

It is important to note that additional steps in the above algorithm may need to be taken when handling longitudinal data. See ref. [10] for more details.

### 7.3 Simulation details

The primary objective in the simulation study is to generate replicate datasets of the observed studies. We use the biological signal from each study as a baseline: both models are fit to estimate the gene expression curves under the alternative and null models. The genes assigned to the alternative model had *q*-values < .1 while genes assigned to the null model had *q*-values *>* .1. The number of unique curves from the alternative model was varied by randomly selecting from the population of genes assigned to the alternative model. For each study and number of unique gene expression curves *g* = 5, 10, 50, 100, 200, the procedure is outlined below:

1. Use the estimated proportion of true nulls 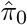 to randomly assign the *m* genes to either the alternative 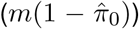 or null 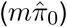 models. Genes assigned to the alternative model followed a unique gene expression curve 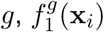. Alternatively, the null genes were randomly sampled from the population of null model fits 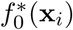
2. Using the observed signal-to-noise ratio (SNR) distribution from the alternative model, calculate an appropriate SNR_M_ such that the estimated number of differential expressed genes at a false discovery rate of 0.1 is close to the observed study. This was done by trial and error: 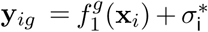, where 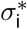 is randomly sampled from the population of standard deviations 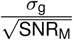 for all *g*.
3. Randomly sample from the population of standard deviations in the previous step to add noise to the alternative model 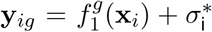 and the null model 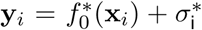 for all genes; call this simulated dataset **Y**^*\**^.
4. Apply the testing procedures to **Y**^*\**^ and calculate *p*-values.
5. Repeat steps (3-4) 500 times and calculate the average number of discoveries and the average false discovery rate for all testing procedures.
6. The estimated proportion of true nulls are shown in Table 1 and the estimated SNR_M_ for the dose, endotoxin, kidney and smoker studies are the 0.86, 0.45, 0.35, and 0.8 quantiles of the SNR distribution, respectively.

### 7.4 True positive enrichment analysis

Consider *i* = 1, 2, *…, m* test statistics *z*_*i*_ calculated on a gene-by-gene basis from a biological study. Given these test statistics, we propose a new summary statistic for gene sets based on the enrichment of true positives. The procedure works as follows. For each gene *i*, we can calculate the local false discovery rate based on the chosen test statistic:

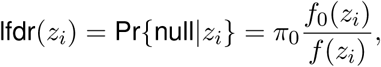

where *π*_0_ is the prior probability that a hypothesis test is null, *f*_0_(*z*_*i*_) is the null density, and *f* (*z*_*i*_) is a mixture of the null and alternative densities [7]. Next, we average the local false discovery rate in gene set *S*,

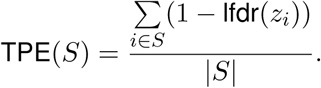

As an example, if we calculated TPE(*S*) = 0.9 then it corresponds to a gene set with an average of 0.9 true positives. Thus this gene set has a high proportion of true positives.

The advantages of working in this framework are (i) it is computationally fast to calculate TPE(*S*) for all gene sets, (ii) the interpretation of important gene sets is more intuitive, and (iii) covariate-adjusted local false discovery rates can easily be incorporated to improve statistical power.

